# *Escherichia coli* Metabolite Profiling Leads to the Development of an RNA Interference Strain for *Caenorhabditis* elegans

**DOI:** 10.1101/731315

**Authors:** Isaiah A. A. Neve, Jessica N. Sowa, Chih-Chun J. Lin, Priya Sivaramakrishnan, Christophe Herman, Youqiong Ye, Leng Han, Meng C. Wang

**Affiliations:** Baylor College of Medicine, Molecular and Human Genetics, Houston, TX, 77030; Baylor College of Medicine, Huffington Center on Aging, Houston, TX, 77030; University of Texas Health Science Center at Houston, Biochemistry and Molecular Biology, Houston, TX, 77030; Howard Hughes Medical Institute, Chevy Chase, MD, 20815

**Author notes:** Corresponding Author: Meng C. Wang, 1 Baylor Plaza N803.03, Houston, TX, 77030. West Chester University, Department of Biology, West Chester, PA, 19383. Cornell University, Department of Ecology & Evolutionary Biology, Ithica, NY, 14850. University of Pennsylvania, School of Medicine, Philadelphia, PA, 19104.

**Keywords:** Metabolomics, microbe-host interaction, metabolism, RNA interference, *C. elegans*

## Abstract

The relationship of genotypes to phenotypes can be modified by environmental inputs. Such crucial environmental inputs include metabolic cues derived from microbes living together with animals. Thus the analysis of genetic effects on animals’ physiology can be confounded by variations in the metabolic profile of microbes. *Caenorhabditis elegans* exposed to distinct bacterial strains and species exhibit phenotypes different at cellular, developmental and behavioral levels. Here we reported metabolomic profiles of three *Escherichia coli* strains, B strain OP50, K-12 strain MG1655, and B-K-12 hybrid strain HB101, and also different mitochondrial and fat storage phenotypes of *C. elegans* exposed to MG1655 and HB101 versus OP50. We found that these metabolic phenotypes of *C. elegans* are not correlated with overall metabolic patterning of bacterial strains, but their specific metabolites. In particular, the fat storage phenotype is traced to the betaine level in different bacterial strains. HT115 is another K-12 *E. coli* strain that is commonly utilized to elicit an RNA interference response, and we showed that *C. elegans* exposed to OP50 and HT115 exhibit differences in mitochondrial morphology and fat storage levels. We thus generated an RNA interference competent OP50 (iOP50) strain that can robustly and consistently knockdown endogenous *C. elegans* genes in different tissues. Together, these studies suggest the importance of specific bacterial metabolites in regulating the host’s physiology, and provide a tool to prevent confounding effects when analyzing genotype-phenotype interactions under different bacterial backgrounds.

## Introduction

Every ecosystem presents its own particular pressures on the plants, animals, and microbes living within. The lives of these macro and microorganisms are often tightly intertwined, and have undergone countless generations of evolution together. The metabolic interaction between microbes and hosts is a crucial aspect of their interplay and is a key factor in shaping metabolic adaptation in the host. In particular, the animal *Caenorhabditis elegans* is known as a powerful model organism for genetic analysis of complex biological processes such as development, behavior, metabolism, and aging. Although long considered as “soil dwelling” nematodes (Brenner, 1974), advances in the field reveal that these animals may be more aptly described as dwelling in “rotting vegetation”(Félix and Braendle, 2010). In this environment, *C. elegans* are exposed to a multitude of bacterial species and strains, many of which can be prey, pathogen, or commensals. These bacteria have different metabolic profiles, contributing to their nutritional inputs, virulence factors, or communication cues, all of which can stimulate different responses in *C. elegans.* The ability of *C. elegans* to detect those metabolic cues derived from bacteria and adjust their physiology accordingly is required for both themselves and their progeny to survive and reproduce in the environment presented to them (Han et al., 2017; Lin and Wang, 2017; Sowa et al., 2015).

Historically, a strain of *Escherichia coli*, known as OP50 has been chosen for growing *C. elegans* in laboratory conditions (Brenner, 1974). OP50 is a uracil-requiring B-type *E. coli* and carries a number of positive traits for *C. elegans* husbandry: it forms a thin bacterial lawn due to its uracil-requirements, it is relatively sticky which facilitates worm transfer, and it is mostly considered non-pathogenic (Couillault and Ewbank, 2002). Alternatively, an *E. coli* B-K-12 hybrid strain HB101 and a ‘wild type’ *E. coli* K-12 strain MG1655 are both healthier than OP50 and can grow into a thicker lawn to support a larger number of *C. elegans* (Lin and Wang, 2017; MacNeil and Walhout, 2013). Furthermore, another *E. coli* K-12 strain, HT115, has been utilized for *C. elegans* gene knockdown using RNA interference (RNAi). HT115 has been rendered capable of housing RNAi inducing double stranded RNA through two major mutations, the loss of the *rnc* allele encoding RNase III, and the introduction of an IPTG inducible T7 polymerase. The phenotypic differences between *C. elegans* grown on OP50 and alternative often K-12 derived strains have been a recent focus of the scientific community. *C. elegans* have been shown to exhibit bacteria dependent molecular and physiological changes, including alterations to development, lifespan, reproductive lifespan, and metabolism (Coolon et al., 2009; MacNeil, et al., 2013; Pang and Curran, 2014; Soukas et al., 2009; Sowa et al., 2015). These phenotypic differences in *C. elegans* are likely associated with metabolic cues derived from different bacterial strains.

In our studies, we have performed high-throughput metabolomic profiling to systemically analyze metabolic difference among three different *E. coli* strains, OP50, HB101 and MG1655. We found that specific bacterial metabolites, rather than gross metabolic patterns, actively influence phenotypic characteristics of *C. elegans* grown on these bacterial strains. Given that *C. elegans* exhibit drastic differences in their mitochondrial morphology, fat storage, and reproductive span when grown on OP50 and HT115, we have generated an OP50 strain that can effectively induce RNAi knockdown and confirmed its efficacy for different tissues and for a variety of endogenous genes. Together the metabolite profiles of different bacterial strains and the OP50 RNAi strain provide useful resources for understanding microbe-host interaction in regulating different physiological activities and for large-scale genetic analysis using RNAi under different bacterial backgrounds.

## Results and Discussion

### Revealing metabolomic diversity among different *E. coli* strains

A microbial community carries diverse species of bacteria, and within each species, there are different strains. Interestingly, *C. elegans* exposed to different strains of *E. coli* exhibit different physiological phenotypes (Brooks et al., 2009; Han et al., 2017; MacNeil,et al., 2013; Sowa et al., 2015), for example, the reproductive lifespan and intestinal fat levels are increased when *C. elegans* are grown on the OP50 *E. coli* strain compared to those animals grown on the HB101 or MG1655 *E. coli* strains (Brooks et al., 2009; Lin and Wang, 2017; Sowa et al., 2015). Using stimulated Raman scattering (SRS) microscopy (Ramachandran et al., 2015; Yong Yu et al., 2014), we quantitatively examined fat content levels in *C. elegans* exposed to those different bacterial strains. In line with previous studies (Brooks et al., 2009), we found that *C. elegans* on OP50 show more fat storage than those on HB101 or MG1655 in the intestine, the major fat storage tissue (Figure 1A). In parallel, we examined mitochondrial fission-fusion dynamics, which have previously been linked to host’s lipid metabolism in response to bacterial inputs (Lin and Wang, 2017). Using a *C. elegans* transgenic reporter strain expressing mitochondrial localized GFP (mito-GFP) under the control of an intestine specific promoter, we imaged mitochondrial morphology and scored the images as one of three distinct categories: filamented, intermediate and fragmented (Figure 1B). Similar to the difference in fat storage, worms grown on OP50 show distinct mitochondrial morphology when compared to those grown on HB101 or MG1655, as evidenced by increased mitochondrial fragmentation in the intestine (Figure 1B).

**Figure 1:**
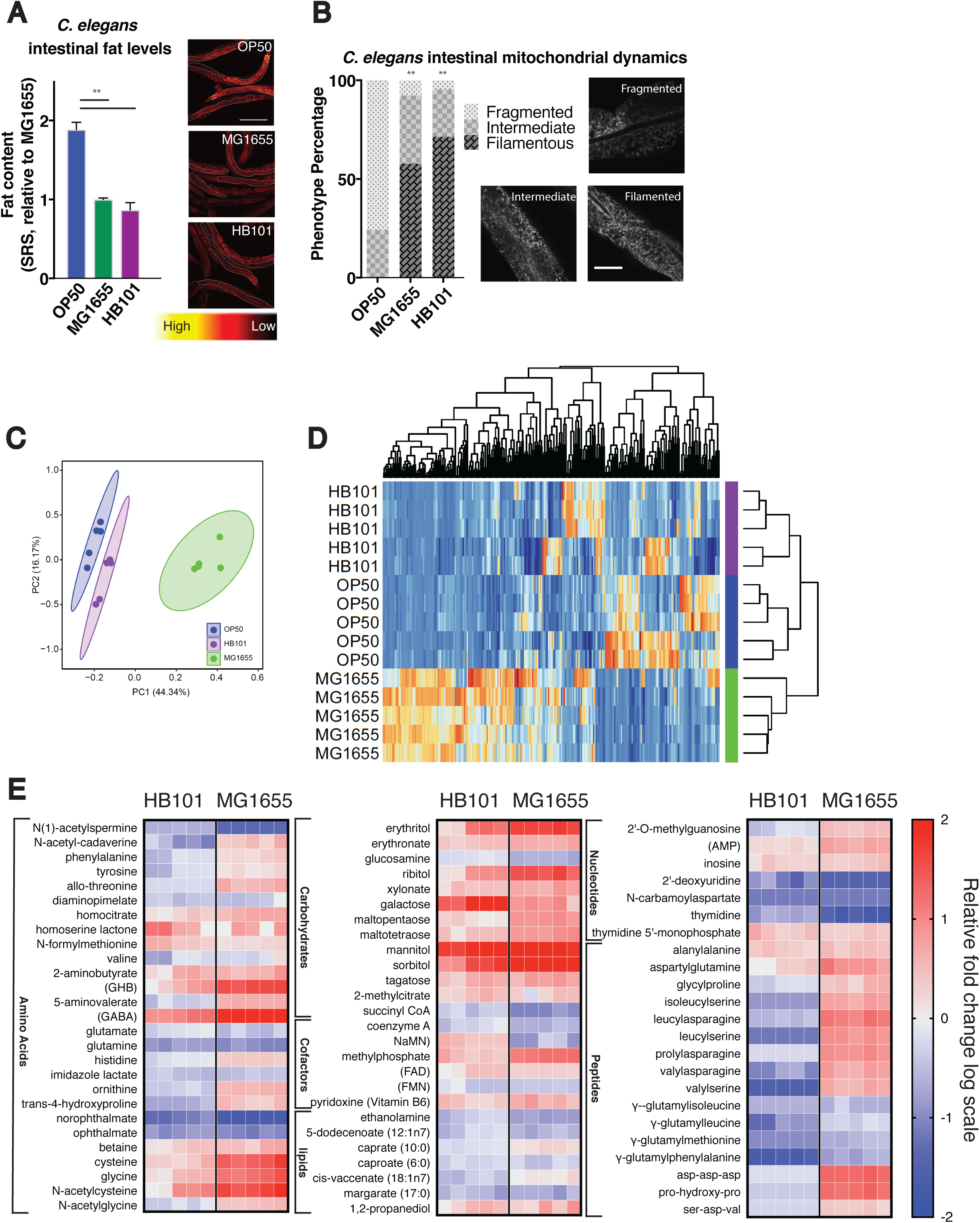
Metabolomic profiles of three *E. coli* strains and their distinct effects on *C. elegans* fat storage and mitochondrial dynamics. **A)** Fat content levels of day-2-old *C. elegans* adults are quantified using SRS microscopy imaging. When compared to those grown on OP50, worms grown on HB101 and MG1655 show decreased fat storage in the intestine. Example SRS images are presented from three independent trials and intestinal areas for quantification are outlined. **B)** Mitochondrial morphology is examined in day-3-old *C. elegans* adults carrying a mitochondria-localized-GFP reporter in the intestine, and classified into three categories, filamented, fragmented or intermediate for quantification. When compared to those grown on OP50, worms grown on HB101 and MG1655 show decreased mitochondrial fragmentation in the intestine. Phenotypes were noted in a double-blind fashion from three independent trials. **C)** PCA analysis of *E. coli* metabolomics profiles shows clustering between OP50 and HB101 that is separated from MG1655. **D)** Clustering of different *E. coli* samples based upon metabolite levels shows that HB101 and OP50 have a similar pattern distinct to MG1655. **E)** Heat maps show relative fold changes in a log scale of significantly altered metabolites in MG1655 and HB101 when compared to OP50. Five independent metabolomics profiles for each strain are presented. Statistical significance indicated by asterisk, * P = <0.05, ** P = <0.005. Error bars represent SEM. Statistical analysis performed using Welch’s two-sample two-sided t-test (E), students t-test (A), or Chi-Squared test (B).

To understand whether the difference in these physiological features of *C. elegans* are associated with the difference in the metabolic features of *E. coli* strains, we systemically analyzed the metabolomic profiles of OP50, HB101, and MG1655 *E. coli* (Supplemental table 1). To our surprise, PCA analysis of all detected metabolites shows that the metabolic profiles of HB101 and OP50 cluster together more readily than OP50 or HB101 do with MG1655 (Figure 1C & 1D). Thus the phenotypic difference of *C. elegans* on different *E. coli* strains is not simply a response to gross metabolic input alterations in those *E. coli* strains, but might be associated with specific metabolites derived from those *E. coli* strains.

We then searched for metabolites that show differences in both HB101 and MG1655 when compared to OP50, which might contribute to the observed *C. elegans* phenotypic differences. Our analysis identified a total of 42 carbohydrate-related metabolites in our bacterial samples (Supplemental table 1), 12 of these metabolites show an increase in both HB101 and MG1655 when compared to OP50 and one shows a decrease (Figure 1E). An inverse relationship is observed in regards to lipid-related metabolites when comparing HB101 and MG1655 to OP50, seven metabolites are differentially observed and six out of the seven exhibit decreased levels in HB101 and MG1655 (Figure 1E, Supplemental table 1). These results suggest that HB101 and MG1655 may have a higher activity of glycolysis and less fermentation than OP50. Among the groups of amino acids and nucleotides, there are also other specific metabolites that show similar changes in HB101 and MG1655, although there is no clear trend as a group.

### Bacterial betaine regulates lipid metabolism in the host

Our previous studies have linked the bacterial one-carbon methyl cycle with lipid metabolism in the host (Lin and Wang, 2017). Among the four metabolites directly involved in the methyl cycle (Figure 2A), methionine is decreased by 50% in HB101 compared to OP50, but not in MG1655, neither dimethyglycine nor homocysteine is significantly changed in HB101 and MG1655, but betaine is increased by 2-fold in HB101 and by 3-fold in MG1655 compared to OP50 (Figure 2B).

**Figure 2:**
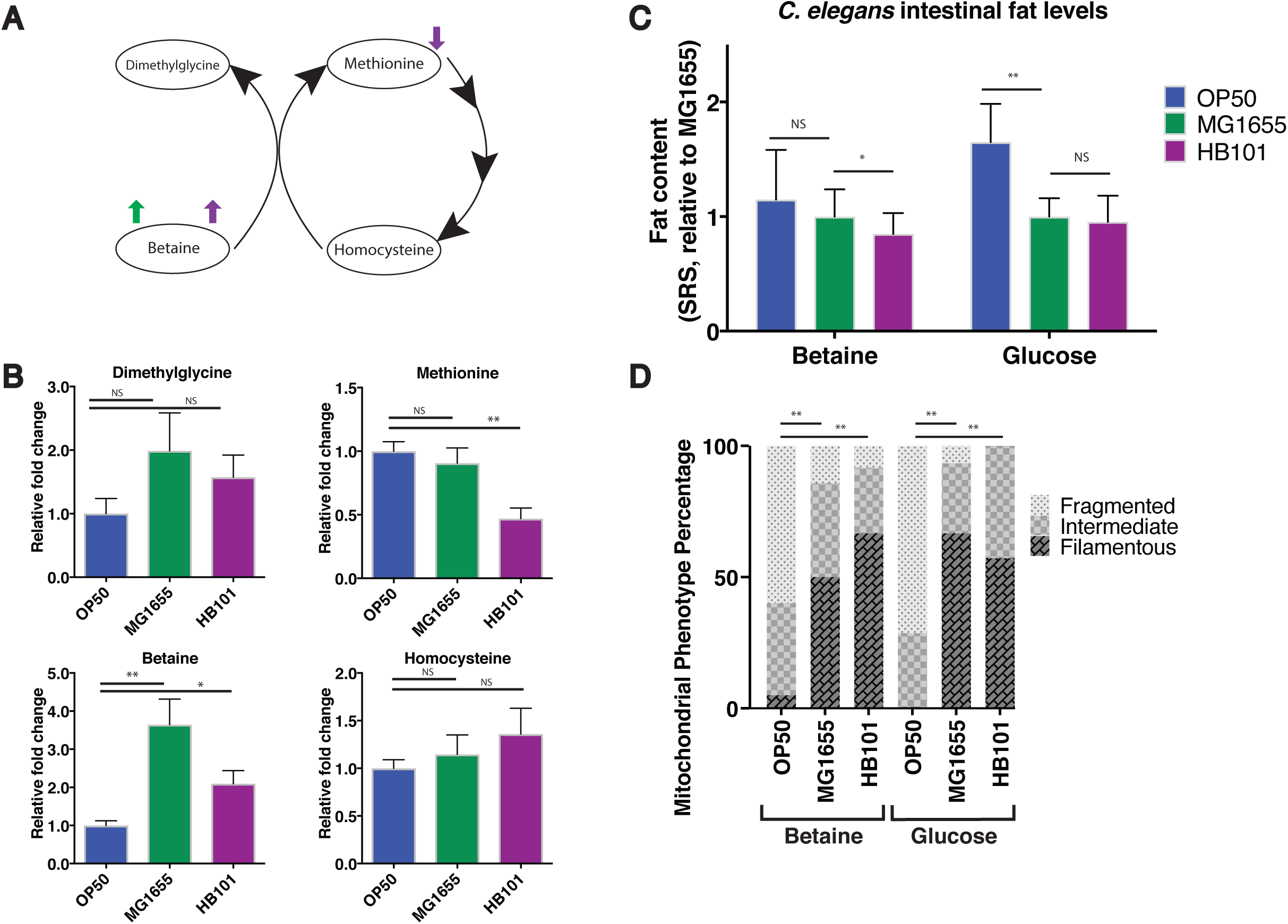
Bacterial metabolite Betaine regulates fat storage in the host. **A)** Simplified one-carbon methyl cycle in *E. coli*, highlighting the methyl donation interactions between four key metabolites, methionine, homocysteine, dimethylglycine, and betaine. Arrows indicate alterations in MG1655 (green) and HB101 (purple) when compared to OP50. **B)** Quantitative comparison of one-carbon methyl cycle metabolite levels show that betaine levels are increased over 3-fold and 2-fold in MG1655 and HB101, relative to OP50, respectively, and the methionine level in HB101 is decreased compared to that in OP50. **C)** Supplementation of betaine but not glucose suppresses the fat storage increase in *C. elegans* grown on OP50. Graphical representation of relative whole intestine fat content levels as measured by SRS microscopy from three independent trials. **D)** Neither betaine nor glucose supplementation affects morphological difference of mitochondria in *C. elegans* grown on HB101, MG1655 and OP50. Graphical representation of mitochondrial dynamics in the intestinal cells from double-blind analyses of three independent trials. Statistical significance indicated by asterisk, * P = <0.05, ** P = <0.005. Error bars represent SEM. Statistical analysis performed using students T-test (B & C), or Chi-Squared test (D).

Next, we examined whether changes in bacterial betaine levels are associated with fat storage differences in the host *C. elegans*. Interestingly, the supplementation of exogenous betaine is sufficient to reduce the high fat storage of *C. elegans* on OP50 to a level comparable to those grown on HB101 or MG1655 (Figure 2C), while betaine supplementation does not further decrease fat storage in worms on HB101 or MG1655. Given the significant induction of carbohydrate-related metabolites in HB101 and MG1655 compared to OP50 (Figure 1E), we also examined whether increased sugar levels contribute to the low fat content levels in *C. elegans* grown on HB101 and MG1655. We supplemented glucose to *C. elegans* grown on different *E. coli* strains, and found that the glucose supplementation is not sufficient to suppress the fat storage difference (Figure 2C). Together, these results suggest betaine as a key metabolite in regulating lipid metabolism in the host *C. elegans*.

We also examined whether betaine regulates mitochondrial fission-fusion dynamics, a trait that has been linked to host’s lipid metabolism in response to bacterial inputs (Lin and Wang, 2017). We supplemented betaine to worms grown on different bacterial strains and examined the effect of this supplementation on mitochondrial dynamics. We found that unlike the fat phenotype, betaine supplementation is not sufficient to suppress the mitochondrial fragmentation phenotype observed in worms grown on OP50 (Figure 2D). In addition, the supplementation of glucose to worms grown on MG1655, HB101 or OP50 also fails to significantly alter mitochondrial dynamics trends when compared the trends observed in non-supplemented animals (Figure 1B). Therefore, betaine can regulate lipid metabolism in the host via a mitochondrial dynamics independent mechanism.

Together the high-throughput metabolite profiles demonstrate that *E. coli* B and K-12 strains exhibit drastic difference in their metabolism, however these global metabolomics patterns of bacteria are not directly associated with their distinct impacts on the physiology of the host. Instead, specific bacterial metabolites contribute to those differences. In the example put forward, bacterial betaine specifically regulates fat content levels in the host *C. elegans*, but has no effect on mitochondrial dynamics. Thus, the association between bacterial metabolism and *C. elegans* physiology is complex and multifaceted, and specific metabolite signals derived from different bacterial strains can be key confounders interfering with the genetic analyses of *C. elegans* phenotypes. In particular, the *E. coli* strain HT115, which is a K-12 strain, has been utilized for over a decade to perform RNAi based screens, verify mutant phenotypes, and generate hypomorphic conditions for otherwise lethal loss of function mutations (Kamath et al., 2001).

### Development of an RNA interference strain using OP50

Almost two decades ago, two large RNAi libraries were generated by Ahringer and Vidal laboratories (Kamath et al., 2003; Rual et al., 2004). Both of these libraries house their RNAi vectors in the HT115 (d3) strain of *E. coli*, which contains a deletion RNase III (*rnc*) allele and an IPTG-inducible T7 RNA polymerase, changes which render the individual bacterium capable of producing and maintaining double stranded RNA from the L4440 double-T7 vector (derived from pPD129.36 (Timmons and Fire, 1998)). Similar to MG1655 in being a K-12 derived strain, HT115 causes reduced fat storage in *C. elegans* when compared to OP50 (Figure 3A), and the reduction level is similar to that caused by either HB101 or MG1655 (Figure 1A). We also examined intestinal mitochondrial morphology and found that worms on HT115 show a more filamented mitochondrial network than that seen in worms on OP50 (Figure 3B). In addition, other physiological characteristics are also different between worms grown on OP50 and on HT115, including reproductive lifespan (Sowa et al., 2015), developmental timing (MacNeil et al., 2013), and lifespan (Pang et al., 2014). Therefore, when comparing phenotypes of *C. elegans* caused by HT115 induced RNAi knockdown, to those exhibited in genetic mutants grown on OP50, the bacterial strain backgrounds introduce an additional confounder that might mislead the interpretation.

**Figure 3:**
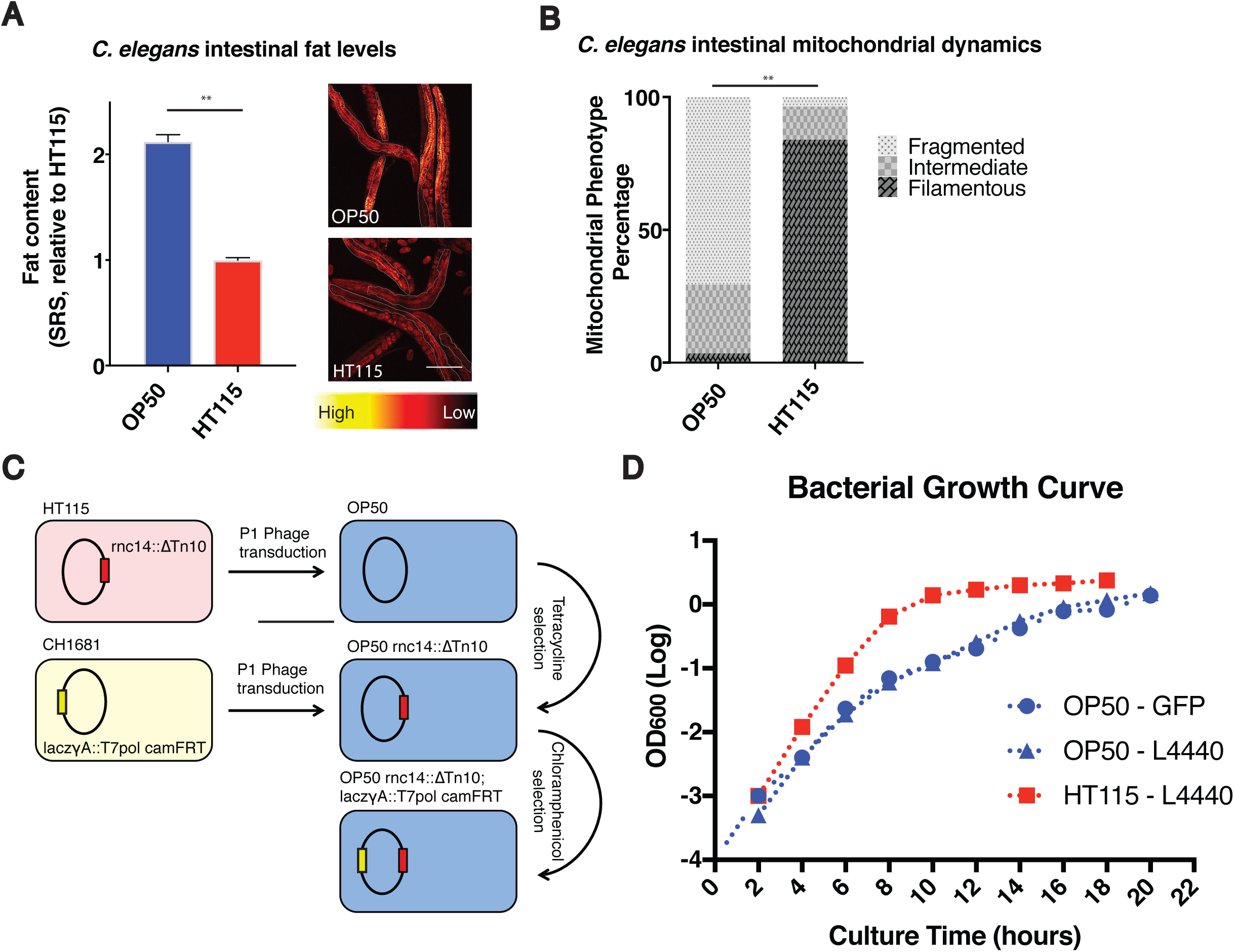
Development of an RNA interference competent OP50 strain. **A)** Quantification based on SRS microscopic imaging shows that worms grown on HT115 have decreased fat storage compared to those on OP50. Example SRS images are presented with intestinal areas outlined for quantification. Graphical quantification is from three independent trials. **B)** Mitochondrial fragmentation, visualized by the intestinal mito-GFP reporter, is reduced in worms grown on HT115 compared to those on OP50. Representative data from two independent trials. **C)** Schematic for the generation of an RNA interference capable OP50 bacterial cell line. **D)** The growth rate of OP50 is lower than that of HT115. Bacterial growth curves for vector carrying OP50 or HT115 show OD_600_ measured every two hours for 20 hours, representative of three independent trials. Statistical significance indicated by asterisk, * P = <0.05, ** P = <0.005. Error bars represent SEM. Statistical analysis performed using students T-test (A), or Chi-Squared test (B).

To override this problem, we have developed an RNAi competent OP50 bacterial strain using phage transduction of two loci required for double stranded RNA production and retention (Figure 3C). The RNAIII RNase (rnc) allele from HT115 (*rnc:14::ΔTn10*) was transduced into the CGC supplied “wild type” OP50 bacteria. This allele provides tetracycline resistance, a trait used for selection of bacterial colonies. Allele introduction was verified using PCR of the appropriate loci (Supplemental figure 1). Following introduction of tetracycline resistance, the *laczγA::T7pol camFRT* allele was introduced by phage transduction to facilitate production of double stranded RNA from the L4440 double T7 vector found in both the Ahringer and Vidal RNAi libraries. This allele is selected for using Chloramphenicol, and its presence was further confirmed using PCR (Supplemental figure1). Following these two phage transduction events, continued rounds of selection on Tetracycline and Chloramphenicol were performed to purify the background of the RNA interference competent OP50 strain (*rnc:14::ΔTn10; laczγA::T7pol camFRT*) herein termed as RNAi competent OP50 or iOP50.

iOP50 *E. coli* carrying the L4440 plasmid show a distinct growth pattern when compared to HT115. We grew iOP50 *E. coli* at 37°C overnight in LB with Carbenicillin (50ug/ml) and measured OD hourly (Figure 3D). We found that a 10 to 14 hour growth period is required for HT115 to enter a stationary phase, while iOP50 requires approximately 18 hours to reach a stationary phase, a growth rate similar to that of non-transformed OP50. Thus, an increased incubation time is necessary for iOP50 strain to provide sufficient, robust and repeatable RNAi knockdown.

### Efficacy of OP50 RNAi strains in gene inactivation

To confirm the knockdown efficacy of iOP50 in different tissues, we first transformed iOP50 with the GFP RNAi plasmid, and supplied those GFP RNAi strains to transgenic *C. elegans* strains expressing GFP in the intestine, muscle, and hypodermis. We found iOP50 induced knockdown of GFP in a qualitatively comparable level to that observed when inducing knockdown using HT115 (Figure 4A). Thus, iOP50-mediated RNAi is sufficient to knockdown genes in different tissues. For these experiments, various growth times have been assessed for iOP50, overnight liquid culture periods ranging from 18 to 22 hours followed by overnight growth on standard NGM plates with 1mM IPTG gives the most robust knockdown (data not shown).

**Figure 4:**
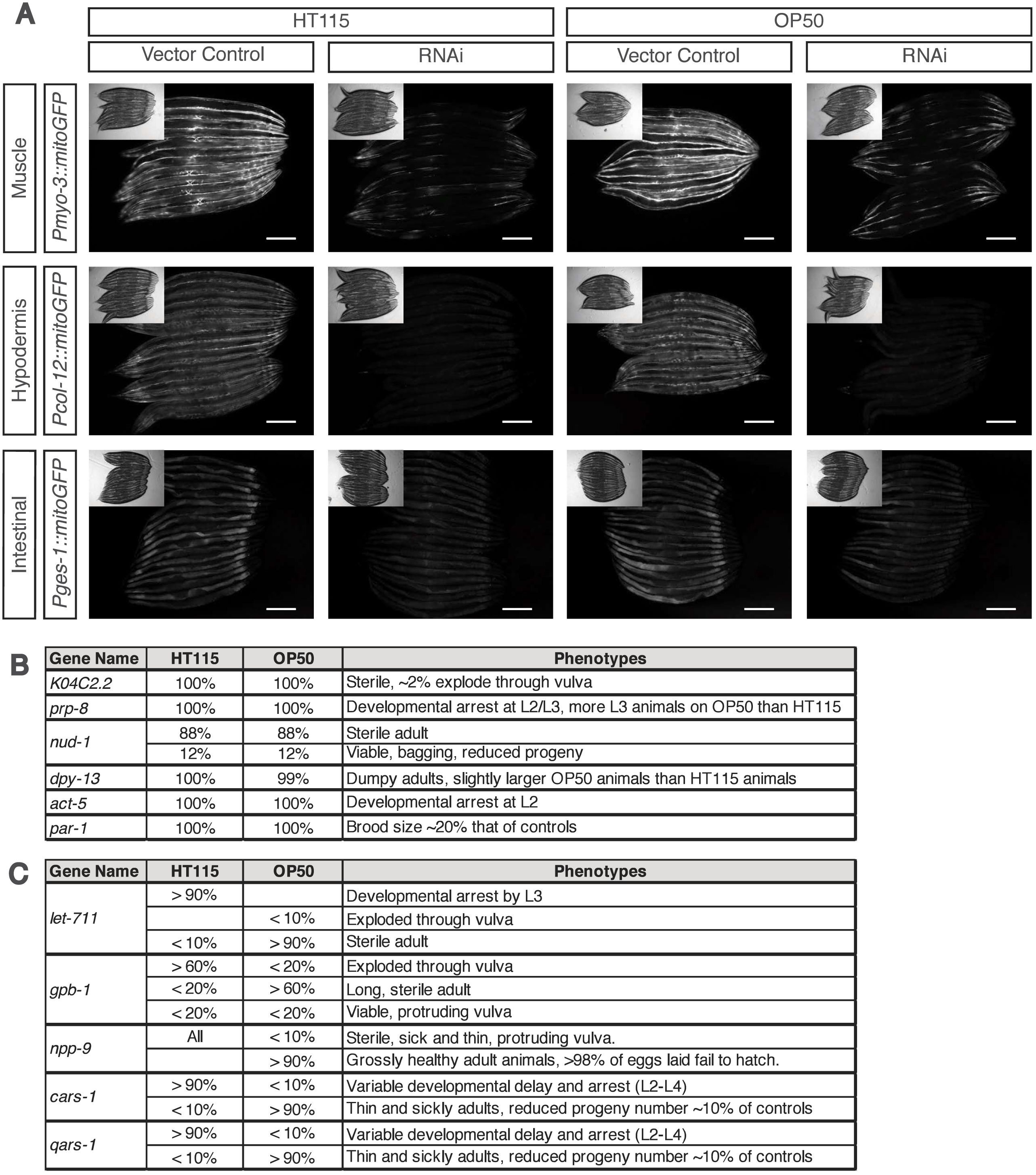
OP50 induced RNA interference is efficient to induce gene knockdown in different tissues. **A)** Example images of GFP knockdown using RNAi in either HT115 or iOP50 background show efficacy in different tissues including the intestine, hypodermis, or muscle. **B)** Gene knockdown using RNAi in either HT115 or iOP50 background gives almost indistinguishable phenotypes. **C)** A number of RNAi lines cause variable phenotype when using HT115 versus iOP50, with less severity using iOP50 than using HT115.

Next, we examined the knockdown efficacy of iOP50 for numerous endogenous genes, where RNAi can induce diverse phenotypes. When comparing the phenotypes of *C. elegans* caused by RNAi knockdown using either iOP50 or HT115, we observed two major categories of results, those where knockdown efficacy is comparable between iOP50 and HT115 (Figure 4B), and those where knockdown using iOP50 gives alternative phenotypes (Figure 4C). The first class includes *K04C2.2, prp-8, nud-1, dpy-13, act-5* and *par-1.* Among them, RNAi knockdown of *prp-8* using HT115 causes developmental arrest by the second larval stage (L2), similarly in iOP50, the majority of animals arrest by L2, although there are few escapers that arrest at the third larval stage (L3); and RNAi knockdown of *dpy-13* leads to decreased body size, but worms are slightly larger when grown on iOP50 than on HT115. The second class includes *let-711, gpb-1, npp-9, cars-1* and *qars-1.* RNAi knockdown of *let-711* using HT115 causes over 90% of the worms to arrest at L3, however when using iOP50, over 90% of worms manage to reach adulthood, but become sterile. For RNAi knockdown of *gpb-1*, the HT115 background generates over 60% dead adults due to explosion through the vulva, but the iOP50 background generates over 60% sterile adults with increased body length. RNAi knockdown of *npp-9* using HT115 gives sterile sick adults, but its knockdown using iOP50 gives grossly healthy adults whose progeny is embryonic lethal. For *cars-1* or *qars-1*, its RNAi inactivation causes developmental arrest when using HT115, but completely penetrant adulthood sickliness and partial adulthood sterility when using iOP50. These phenotypic differences between HT115 and iOP50 RNAi might be related to the strength of gene knockdown, which may be weaker with iOP50, and might also be related to the specific role of certain *C. elegans* genes in response to different bacterial strains. Together, these studies demonstrate the efficacy of iOP50 in executing RNAi knockdown in different tissues and for various genes, and also highlight the importance of examining the effect of *C. elegans* genes under different bacterial strain backgrounds.

## Conclusion

In summary, *C. elegans* exhibit drastically different phenotypes when exposed to different bacterial strains, which are unrelated to genetic alterations in *C. elegans.* These differences could introduce confounders that complicate analysis of genetic regulation, but also provide researchers the opportunity to investigate environment-microbe-host interactions. Our metabolomics studies systematically reveal metabolite difference among different *E. coli* strains, and found that *C. elegans* phenotypic changes are directly associated with specific bacterial metabolites. In particular, we have linked betaine, a metabolite derived from the methyl cycle, with lipid metabolism in worms, which however does not contribute to mitochondrial dynamics or reproductive span (Supplemental figure 2). This one-to-one relationship between a microbial metabolite and a host’s phenotype highlights the importance of investigating the mechanistic impact of microbial metabolism on host’s physiology beyond profiling microbial phylogenetic composition. Moreover, to expand the analysis toolkit of genetic regulation and microbe-host interaction, we have rendered the common *E. coli* strain OP50 competent for the induction of the RNAi response in *C. elegans*. This bacterial strain contains the same two alleles that render HT115 capable of inducing the RNAi response. These changes allow iOP50 to induce RNAi in multiple tissues to a level similar to that found in HT115 fed animals. A similar OP50 RNAi strain has also been generated and used for lifespan screens (Xiao et al., 2015). iOP50 has recently been used in our lab as the basis for a small scale RNAi screen (96 wells) and has proven amenable to en-mass transformation. Phenotypes observed and described in the iOP50 screen hold true for the reciprocal HT115 screen (data not shown). The strain is available by request, and has been deposited with the CGC for academic use. If your mutant *C. elegans* strain fails to demonstrate the same phenotype as your RNAi fed animals or vice-versa, we implore you to pause and question whether the bacterial strain is responsible for the discrepancy and apply iOP50 into your phenotypic validation.

## Supporting information

Supplemental table 1 - Metabolites

---

**Supplemental Table 1 –** Metabolite levels for five samples of each strain of OP50, HB101, and MG1655. Levels are normalized to total pellet weight and Bradford protein.

**Supplemental Figure 1.**
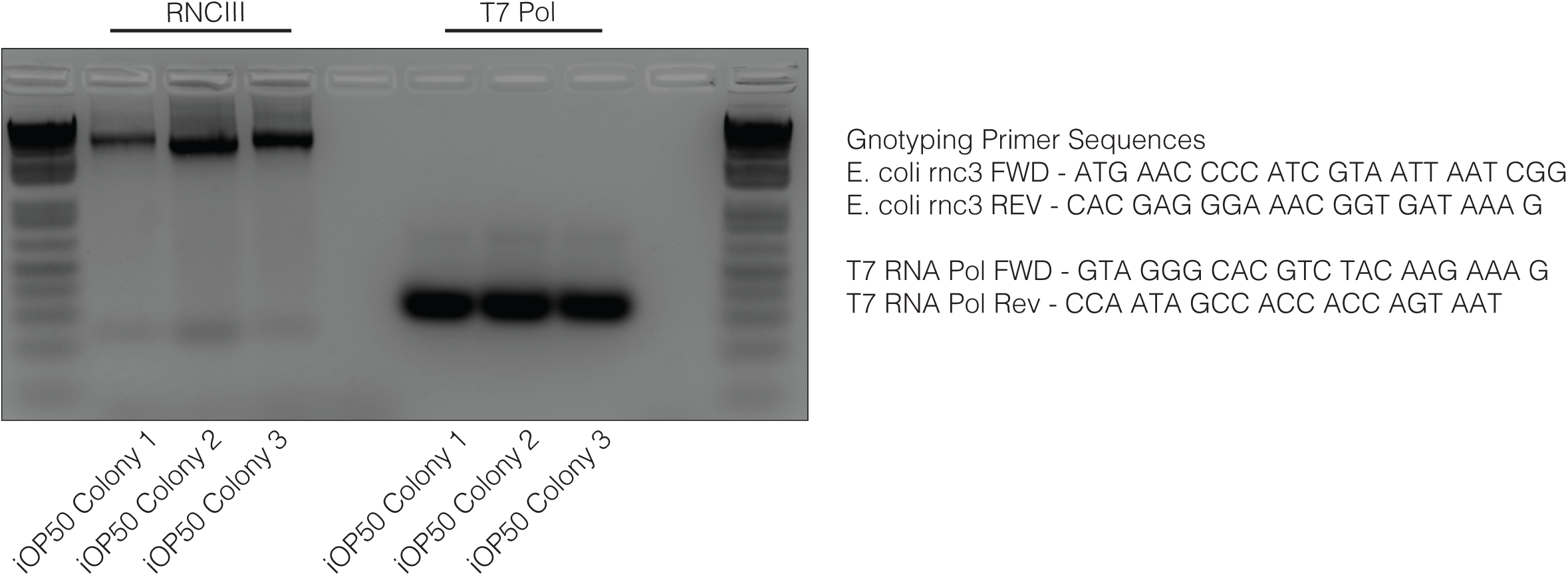
2% agarose gel depicting three colonies of iOP50 with the rnc deletion allele amplified, and the same three colonies of iOP50 with the T7 RNA polymerase allele identified.

**Supplemental Figure 2.**
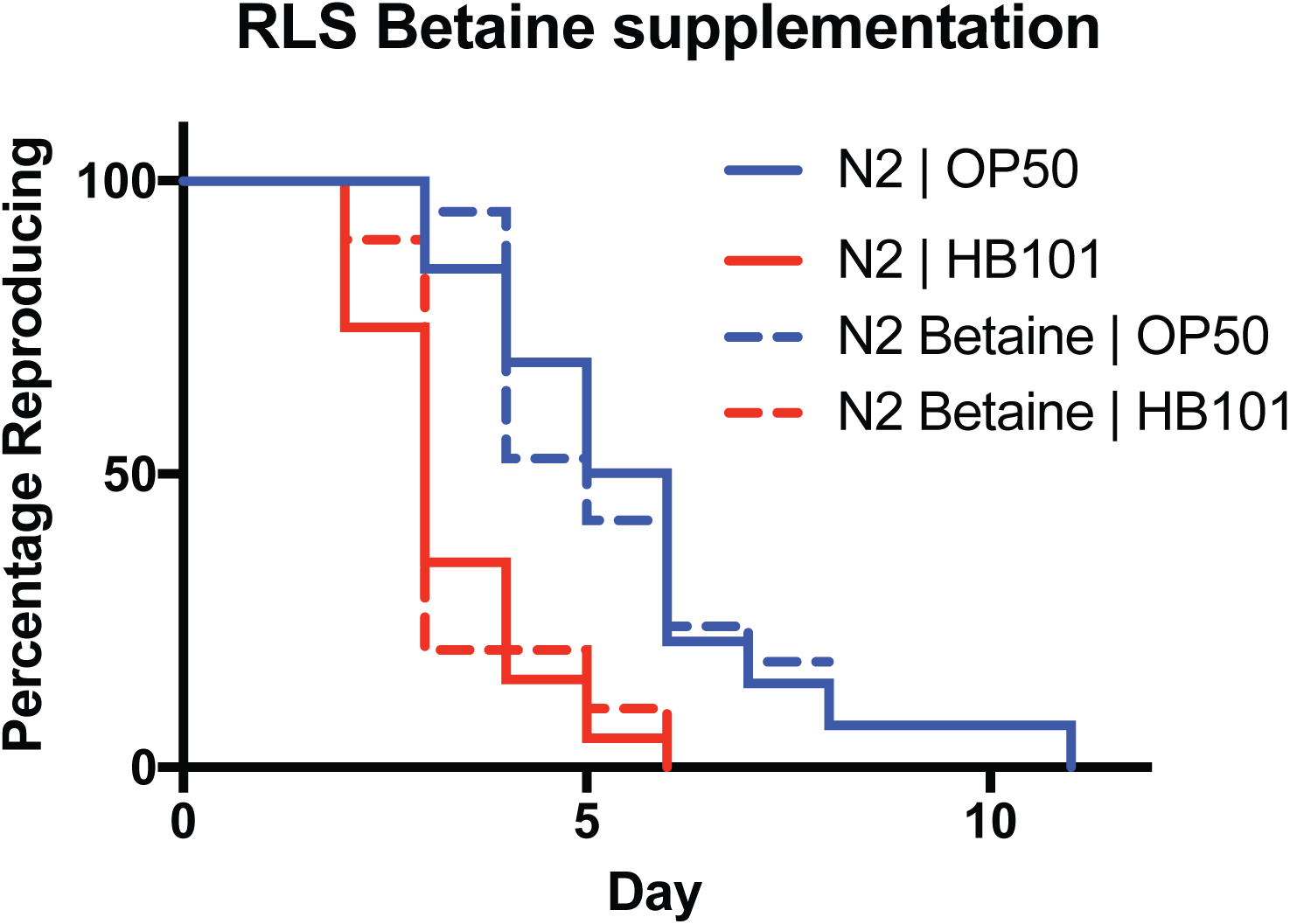
Reproductive lifespan of animals grown on OP50 or HB101 and supplemented with Betaine or vehicle controls.

## Materials & Methods

### *C. elegans* strains used

The *C. elegans* strains used in this study were either provided by the Caenorhabditis Genetics Center (CGC), which is funded by NIH Office of Research Infrastructure Programs (P40 OD010440), or made in house by gonadal microinjection of DNA mixtures at the young adult stage. Integration of extrachromosomal arrays were induced by gamma irradiation exposures (4,500 rad for 5.9min) at the L4 stage, and the integrated progeny were backcrossed to N2 at least five times. Worm strains: N2 (Bristol), *raxIs46[Pges-1::sl2::gfp], raxIs49[Pcol-12::mitogfp], raxIs51[Pges-1::mitoGFP], raxIs52[Pmyo-3::mitogfp]*.

### Bacterial strains used

The following bacterial strains were used in this study. *E. coli* wild type (MG1655) was a gift from J.J. Wang, and CH1681 was a gift from C. Herman. The *E. coli* strains OP50, and HB101 were obtained from the CGC. The *E. coli* strain HT115(d3) was obtained from the Ahringer (Kamath et al., 2003) RNAi library.

### Generation of RNA interference (RNAi) competent OP50 bacteria

Standard OP50 (CGC) was rendered RNAi competent through two modifications facilitated by phage transduction. The OP50 RNAIII RNase (*rnc*) allele was replaced with the deletion allele *(rnc14::ΔTv10)* found in HT115(D3). Subsequently a genomically encoded IPTG-inducible T7 RNA polymerase *(laczγA::T7pol camFRT)* was introduced into OP50*(rnc14::ΔTv10)* from CH1681. In brief, OP50 overnight cultures were transduced with HT115*(rnc14:: ΔTv10)* lysate, insertion events were selected for using tetracycline (Tet) resistance. Individual Tet resistant colonies were grown in LB + Tet (50ug/ml) and banked, PCR was then used to confirm the insertion of the *rnc14::ΔTv10* allele by PCR. OP50(*rnc14::ΔTv10*) overnight culture was then transduced with CH1681 (*laczγA::T7pol camFRT*) P1 lysate. Insertion events were selected for using Chloramphenicol (CAM, 17ug/ml). Three rounds of population expansion and bottlenecking using Tet + CAM (50ug/ml, and 17ug/ml) selection was used to purify the RNAi inducible OP50 (iOP50) background. Individual colonies were banked, and PCR was used to confirm the presence of *rnc14::ΔTv10* and *laczγA::T7pol camFRT* (Supplemental figure 1). iOP50 was then transformed with L4440 plasmids targeting GFP, and RNAi efficacy was determined in *C. elegans* strain *raxIs46[Pges-1::sl2::gfp].*

### Producing chemically competent iOP50 bacteria

All steps are performed using sterile technique. Bacterial iOP50 stocks from −80°C were streaked onto Tetracycline (Tet at 50ug/ml) and Chloramphenicol (CAM at 17ug/ml) containing LB and grown overnight at 37°C. Two tubes of 2ml LB with Tet and CAM (50ug/ml and 17ug/ml) were inoculated with single colonies of iOP50 and grown overnight in a 37°C shaker. The next day, 1ml of overnight culture is used to inoculate 100ml of LB without antibiotics. The culture is shaken at 37°C for 1.5 to 3 hours, until the OD600 is between 0.2 and 0.5. During the shaking, sterile microcentrifuge tubes (>50), 50ml centrifuge tubes, 100mM CaCl_2_, and 100mM CaCl_2_/15% Glycerol are cooled to 4°C. The bacteria are then transferred to two 50ml cold centrifuge tubes. Chill the cells on ice for 10 minutes, all future steps are conducted in the 4°C cold room, in a pre-chilled 4°C centrifuge or with the bacteria kept on ice. Harvest the cells by centrifugation at 4000rmp for 3 minutes, then remove and discard the supernatant. Cells are then gently re-suspended in 5ml of cold 100mM CaCl_2_. Incubate the cells on ice for 20 minutes and then harvest the cells by centrifugation at 4000rmp for 3 minutes. Return the cells to ice and discard the supernatant. Gently re-suspend the cells in 2ml of cold 100mM CaCl_2_/15% Glycerol and then dispense into microcentrifuge tubes in 50µl aliquots. Store the cells at −80°C. Thaw the cells on ice and transform with custom plasmids, or those obtained from the Ahringer or Vidal library (Kamath et al., 2003; Rual et al., 2004).

### Transformation of iOP50

Competent cells were transformed using standard methodologies described in the New England Biotechnology protocol (C2987H/C2987I).

### Metabolomics analysis

*E. coli* metabolite profiling was performed at Metabolon, Inc (Durham, NC) with 5 biological replicates of each test bacterial strain (MG1655, OP50, HB101). *E. coli* colonies were inoculated in LB, grown overnight, and then plated on Nematode Growth Medium (NGM) Agar plates to form lawns for three days at 20°C. Lawns were then scraped off of the surface of the agar and collected in distilled deionized water, samples were then washed 3 times. Each sample contained approximately 0.1mg of pooled bacterial pellet which was then flash frozen using liquid nitrogen. Metabolomics profiling was performed in the method described previously (Evans et al., 2009). Statistical analysis was performed using Welch’s two-sample t-test to identify compounds that differed significantly between bacterial strains.

### Measurement of lipid content levels in *C. elegans* intestine by SRS microscopy

*C. elegans* () were raised from L1 on the specified food source/supplement (Betaine at 200mM or Glucose at 0.4% final volume compared to agarose). Day-2-old adults (∼10 for each of three biological replicates) were anesthetized in 1% sodium azide in M9 buffer (3g KH2PO4, 6g Na2HPO4, 5g NaCl, 1ml 1M MgSO4, H2O to 1 liter, sterilized by autoclaving), and mounted on 2% agarose pads sandwiched between glass microscopic slides and coverslips. Images were taken using the SRS system as described in previous studies (Ramachandran et al., 2015; Wang et al., 2011). In brief, when using Stokes beam at 1064nm and pump beam at 817nm, the energy difference between the Stokes and pump photons resonates with the vibrational frequency of CH2 bonds (2845cm-1). Due to the fact that the CH2 chemical bonds are highly enriched in lipid carbon chains, we quantitatively image total fat content levels by detecting the emitted Raman signals of CH2 bonds.

### Mitochondrial morphology observation and dynamics measurement

The intestinal mitochondrial morphology was examined using day-3-old adult transgenic *C. elegans* strain *raxIs51[Pges-1::mitoGFP]* raised from L1 on the specified food source, with the specified supplement (Betaine at 200mM or Glucose at 0.4% final volume compared to agarose). Images are presented as single-layer con-focal images in anterior regions for consistency. For sample preparation, worms were anesthetized in 1% sodium azide (NaAz) in M9 buffer, and mounted on 2% agarose pads sandwiched between glass microscopic slides and coverslips. Confocal images were taken using an IX81 microscope (Olympus) connected to an AxioCam ICc3 camera (Zeiss). For morphological categorization, if the lengths of majority of mitochondrial filaments in a cell were longer than ∼4µm, it would be considered “filamented”; if the lengths of majority of mitochondrial filaments in a cell were shorter than ∼2µm, it would be considered “fragmented”; the rest belonged to the category “intermediate”. Categorical rankings were determined by three blinded scorers, rankings were only determined for cells on which at least 2 out of 3 of the scorers agreed. An individual cell was utilized as the unit, at least three trials were performed and the accumulated counts from each genotype and environmental condition were summed up across biological replicates, percentages were calculated and reported as bar representations. The chi-squared test for trend was conducted for statistical analysis.

### Inducing RNAi using HT115 or iPO50

HT115 or iOP50 bacterial cultures were grown in LB with Carbenicillin (2.64µM) for 14h, and 18h respectively. 150µl of bacterial culture was then plated on RNAi plates (standard nematode growth medium (NGM) agarose plates with IPTG (1mM) & Carbenicillin (2.64µM)) and allowed to dry. Production of dsRNA was induced overnight (18h) at room temperature. *C. elegans* were then added to the plates. All noted phenotypes or images were taken in P0 animals grown 62-72h at 20°C.

### Florescent imaging for determining RNAi efficacy

Well-fed N2 animals were raised for two generations on standard NGM plates seeded with OP50. Animals were synchronized using a variation of the “egg prep” methodology described in (Porta-de-la-Riva et al., 2012). ∼100 animals were grown from the L1 stage on RNAi plates with bacteria containing L4440 based plasmids targeting GFP. At 72h post plating, ∼10 day 1 adult animals were picked into a pool of 8µl 2% sodium azide in M9 buffer located in the center of a freshly made empty NGM plate. Animals were paralyzed, and then arranged by gentle prodding and by increasing the surface area of the NaAz pool, while allowing it to evaporate. Plates were then immediately imaged using a Hamamatsu digital camera (C11440), utilizing the x-cite xylis light source, and a Nikon SM218 microscope.

### Determining RNAi induced phenotypic alterations

Well-fed N2 animals were raised for two generations on standard NGM plates seeded with OP50. Animals were synchronized using a variation of the “egg prep” methodology described in (Porta-de-la-Riva et al., 2012). 6cm plates of bacteria containing L4440 based plasmids targeting the loci of interest from the Ahringer or Vidal library were prepared. ∼100 animals were grown from the L1 stage on each of the 6cm RNAi plates. Five independent trials were conducted and results were collated. Animals were grown for ∼72h at 20°C, phenotypes were then noted for all animals on the plates and penetrance was determined.

### Methodology for performing Reproductive Lifespans

Reproductive lifespans (RLS) were performed at 20°C on animals that were developmentally synchronized using a variation of the “egg prep” methodology described in (Porta-de-la-Riva et al., 2012). Well-fed animals are raised for two generations on standard Nematode Growth Medium (NGM) plates seeded with OP50. The third generation of animals are plated on antibiotic free NGM plates seeded with the test bacteria, and if necessary, an inducible agent (IPTG) or supplement (Betaine at 20mM during development, and 100mM after L4, based upon plate volume). Upon development to the L4 stage, worms are singled to individual plates containing their specified food source, the animals are then transferred to fresh plates every day at the same time until reproduction cessation, determined when no progeny are produced for three continuous days. Two days after the individual worms are transferred, plates are checked and then double checked for progeny. Animals which bag or die are noted as censors. Reproductive span curves were calculated using Kaplan-Meier survival analysis and compared using the log-rank test. At least two independent trials were conducted.

## Author Contributions

I.A.A.N, J.N.S., C.J.L. and M.C.W. conceived the study, I.A.A.N., J.N.S., and C.J.L. conducted experiments and analysis. Y.Y. and L.H. conducted clustering analysis. J.N.S, P.S. and C.H. created the iOP50 line. I.A.A.N. and M.C.W. wrote the manuscript.

## Acknowledgements

We thank M. Park and P. Hu for assistance with blinding and ranking mitochondrial dynamics images. A. Dervisefendic & P. Hu for experimental support. C. Herman & J. Wang for providing bacterial lines. This work is supported by: NIH grants R01AT009050, DP1DK113644, R01AG062257, R01AG045183 (to M.C.W.), Howard Hughes Medical Investigator grant (to M.C.W.), March of dimes foundation grant #l-FY16-244 (to M.C.W.), Burroughs Wellcome Foundation Houston Laboratory and Population Sciences Training Program in Gene-Environment Interaction (to J.N.S.), NIH grant R01GM088653 (to C.H.), and the Cancer Prevention and Research Institute of Texas (grant no. RR150085) to CPRIT Scholar in Cancer Research (L.H.).

